# Polymer-assisted intratumoral delivery of ethanol: Preclinical investigation of safety and efficacy in a murine breast cancer model

**DOI:** 10.1101/2020.05.29.123125

**Authors:** Corrine A. Nief, Robert Morhard, Erika Chelales, Daniel Adrianzen Alvarez, Ioanna Bourla, Christopher T. Lam, Alan A. Sag, Brian T. Crouch, Jenna L. Mueller, David Katz, Mark W. Dewhirst, Jeffrey I. Everitt, Nirmala Ramanujam

**Author notes:** Corresponding author: Corrine Audrey Nief; 101 Science Drive, Box 90281, Durham, NC 27708.

## Abstract

Focal tumor ablation with ethanol could provide benefits in low-resource settings because of its low overall cost, minimal imaging technology requirements, and acceptable clinical outcomes. Unfortunately, ethanol ablation is not commonly utilized because of a lack of predictability of the ablation zone, caused by inefficient retention of ethanol at the injection site. To create a predictable zone of ablation, we have developed a polymer-assisted ablation method using ethyl cellulose (EC) mixed with ethanol. EC is ethanol-soluble and water-insoluble, allowing for EC-ethanol to be injected as a liquid and precipitate into a solid, occluding the leakage of ethanol upon contact with tissue. The aims of this study were to compare the 1) safety, 2) release kinetics, 3) spatial distribution, 4) necrotic volume, and 5) overall survival of EC-ethanol to conventional ethanol ablation in a murine breast tumor model. Non-target tissue damage was monitored through localized adverse events recording, ethanol release kinetics with Raman spectroscopy, injectate distribution with *in vivo* imaging, target-tissue necrosis with NADH-diaphorase staining, and overall survival by proxy of tumor growth. EC-ethanol exhibited decreased localized adverse events, a slowing of the release rate of ethanol, more compact injection zones, 5-fold increase in target-tissue necrosis, and longer overall survival rates compared to the same volume of pure ethanol. A single 150 µL dose of 6% EC-ethanol achieved a similar survival probability rates to six daily 50 µL doses of pure ethanol used to simulate a slow-release of ethanol over 6 days. Taken together, these results demonstrate that EC-ethanol is safer and more effective than ethanol alone for ablating tumors.

**Graphical Abstract:** The inclusion of ethylcellulose limits extra-tumoral leakage of ethanol and increases the target-tissue ablation.

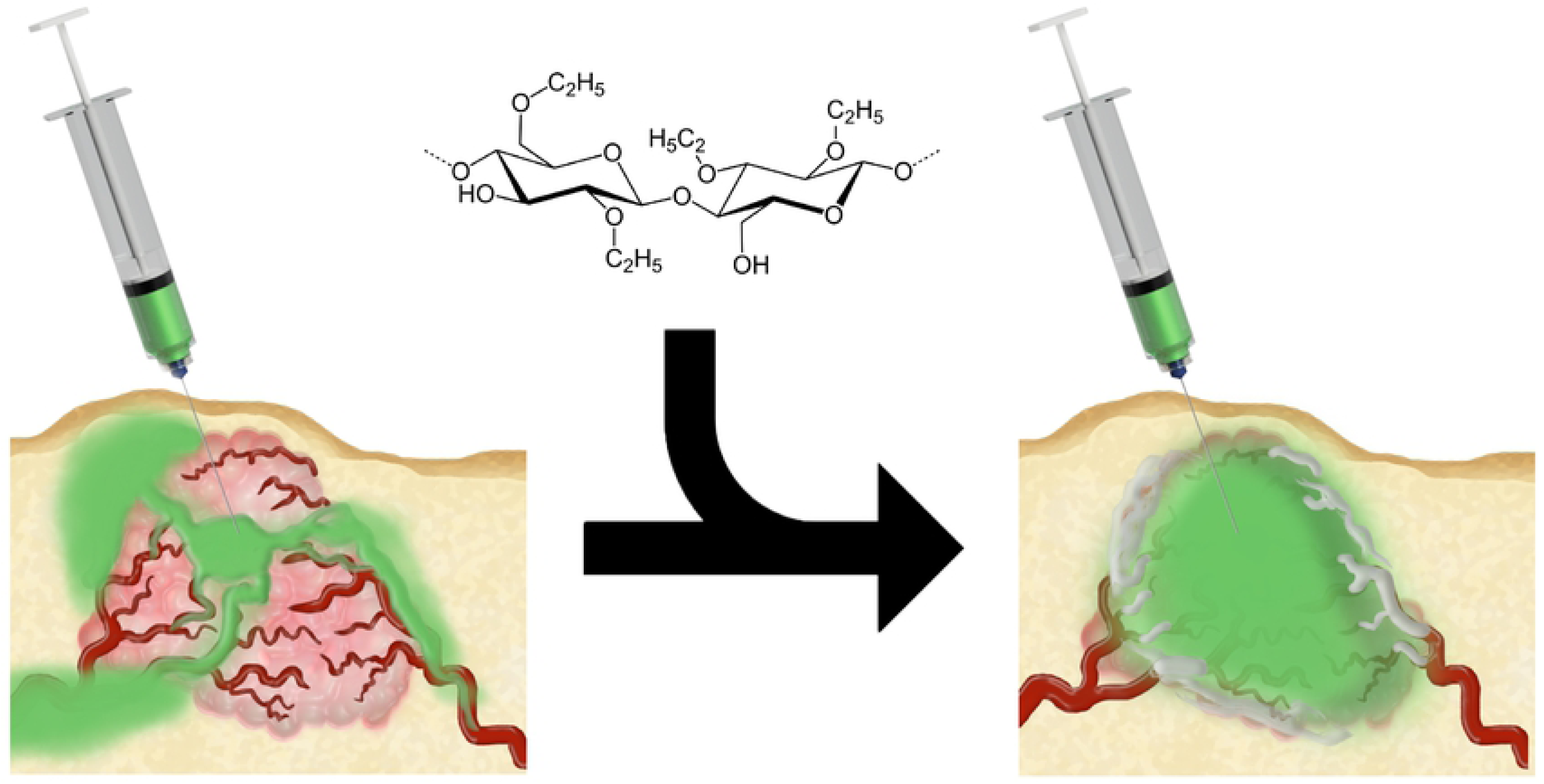

## Introduction

Ablation is the focal destruction of tissue using a small instrument delivered under the skin, typically performed when surgical excision of the tissue is impractical, inaccessible, or dangerous for the patient. Ethanol ablation, or percutaneous ethanol injection (PEI), kills tumors by causing coagulative necrosis upon contact with tissue (*1-3*). Ablation of small focal tumors with ethanol is a well-accepted technique because it is cost-effective, can be visualized with ultrasound, and has acceptable clinical outcomes for treatment of hepatocellular carcinoma (*4*). Ethanol ablation has been used as an alternative to radiofrequency ablation for hepatocellular carcinomas in cirrhotic patients because it is less expensive and less time-consuming with a similar 5-year survival rate for small lesions (*3, 5*). More recently, ethanol ablation has been used successfully for neurolysis (*6, 7*), cardiomyopathy septal ablation (*8, 9*), treatment of cystic thyroid nodules (*1, 2, 10, 11*), palliation of osteolytic bone metastases (*1, 2, 12, 13*) and ablation of neuroendocrine tumors (*14, 15*). Notably, the precision with which any ablation modality can inflict necrosis on a target tissue without damaging non-target tissue is the key metric for success of an ablation technique. Unfortunately, pure ethanol leaks freely out of the tumor, along the path of least resistance resulting in high rates of post-treatment tumor progression (*16, 17*). In fact, ethanol ablation often requires multiple sessions to achieve the same efficacy as radiofrequency ablation (*18*) as it provides incomplete coverage of large lesions in a single session (*19*). For these reasons, microwave ablation and cryoablation have gained favor in high-resource settings, as tissue destruction can be directed to a radially symmetric volume of tissue more reliably. However, use of thermal ablation requires hard to access resources including CO_2_ cylinders for cryotherapy, expensive specialized machinery for microwave ablation, and a consistent power supply for all thermal ablation limiting use of thermal ablation to high-resource settings. Therefore, an unmet clinical need remains for local treatment, especially in low-resource settings, where surgery remains widely inaccessible (*20*).

Here a simple method for percutaneous tumor ablation using polymer-assisted delivery of ethanol with ethyl cellulose (EC) is demonstrated in murine tumors. EC is an inert, ethanol-soluble polysaccharide used as a coating for medical pills and regarded as safe by the Food and Drug Administration. Once the EC-ethanol mixture encounters an aqueous environment, a fibrous gel is formed via non-solvent induced phase separation when water contacts the EC-ethanol mixture (*21*). EC was incorporated into the ethanol solution in order to increase the viscosity of the ethanol and limit the spread of ethanol out of the tumor. We hypothesize that EC will increase the time that ethanol is held at the injection site in an intratumoral depot, and as the ethanol remains in contact with the target-tissue longer it will increase local necrosis. In previous studies we characterized the rheological properties of the gel produced by EC-ethanol (*22*) and its delivery into *ex vivo* tissue (*23*). We previously demonstrated that 3% EC-ethanol forms a fibrous gel upon contact with water and in superficial hamster cheek pouch tumors decreased tumor volume observed over a 7 day period compared to pure ethanol (*22*). EC-ethanol’s unique physical properties have been safely utilized clinically to treat venous malformations (*24, 25*) and herniated discs (*26*), but has yet to be assessed for the ablation of subcutaneous tumors.

In this study we utilized murine breast cancer tumors to test the ability of EC-ethanol to precisely ablate tumors percutaneously. Cell death due to ethanol is dependent on two factors: the concentration of ethanol and the exposure time. Thus, our goal was to slow the release of ethanol from the injection site to allow for maximum ethanol exposure to the tissue near the needle while limiting ethanol exposure to non-target tissue. Ethanol’s cytotoxic ability is due to the fact ethanol molecules are very-small and polar; thus, tagging ethanol with a radio-opaque or fluorescent probe to monitor its location was deemed impractical for this study as it would affect the transport of ethanol through tissues. Ethanol does, however, produce a distinct signal with Raman spectroscopy (*27, 28*) which was utilized to monitor the release kinetics of ethanol through tumor tissue. To monitor the spatial distribution of EC-ethanol a fluorescent powder, fluorescein, was mixed with EC-ethanol to monitor the approximate location of the injectate *in vivo* and *ex vivo*. Fluorescein acts as a representative small molecule, to demonstrate how chemotherapeutic agents could easily be incorporated into the EC-ethanol solution to improve intratumoral delivery. Ablation efficacy was then validated with viability staining of tumors and monitoring the tumor volume after EC-ethanol ablation.

Large 1 cm^3^ mouse tumors were used to model clinically relevant tumor sizes for assessment of fluid distribution (fluorescein distribution) and tumor necrosis (viability staining). To assess safety (adverse events) and efficacy (tumor growth) small 50 mm^3^ tumors were used to allow for sufficient time between the treatment and the ethical tumor burden limit of 2000 mm^3^ in mice. 67NR cells were selected for this study because they are non-metastatic and grow rapidly without producing large necrotic cores, unlike their sister line 4T1, which could artificially contain injection fluids. 67NR cells are triple-negative breast cancer with a luminal phenotype and BALB/c background. This study characterizes how the inclusion of EC alters ethanol delivery and the resulting safety and efficacy of percutaneous EC-ethanol injections in subcutaneous breast tumors.

## Materials and Methods

### 67NR Cell culture and Tumor Implantation in Mice

Subcutaneous flank tumors were established in 6-8-week-old female nu/nu mice (Duke University CCIF) or BALB/c mice (Charles River Labs) through injection of 5×10^5^ 67NR murine luminal breast cancer cells in 100 μL of serum-free RPMI (VWR). 67NR cells are locally invasive but non-metastatic. Murine breast cell line 67NR was provided by Dr. Fred Miller (Karmanos Cancer Institute, Detroit, MI) through Dr. Inna Serganova and Dr. Jason Koucher (Memorial Sloan Kettering Cancer Center, New York, NY). 67NR cells were cultured in RPMI with 5% penicillin-streptomycin. Mice were euthanized when tumors reached 1500 mm^3^ or body weight dropped 15% below baseline. Tumors were excised post-euthanasia, flash frozen, and processed by the Duke Substrate Core (Duke University, Durham, NC).

### EC-ethanol Injections

Anhydrous ethanol was obtained from Koptetc (King of Prussia, NJ). Ethyl cellulose (99% purity, 100 cP) was purchased from Sigma Aldrich. Fluorescein (free acid, 95% purity) was purchased from Sigma Aldrich. All experiments used 2.5% w/w of fluorescein mixed in ethanol. The stability of EC polymer is well documented (*29*), however the rate of EC degradation in pure ethanol has yet to be quantified. To limit possibility of chemical modification of EC polymers, 6% EC-ethanol solution was mixed without heat no more than 24 h before injection. All mixtures were mixed with an ethanol safe stir-bar in ethanol-safe containers. All injections were performed using 27-gauge needles with manual needle placement in the center of the tumor. Injections were performed at a fixed rate of 1 mL/h with a syringe pump unless otherwise specified.

### Adverse Events Recording in Mice

All animal work was performed under protocols approved by the Duke University Institutional Animal Care and Use Committee. All experiments were performed in accordance with relevant animal welfare guidelines and regulations (Protocol A160-18-07). Murine 67NR tumors were grown to 5-10 mm diameter in the flank of 116 BALB/c or nu/nu mice. EC-ethanol injection volumes ranged between 2-16 mL/kg. For comparison, 16 mice were given a single intratumoral injection of 6 mL/kg of ethanol and fluorescein alone, 5 mice received 50 µL of ethanol daily for 6 days referred to as the “repeated ethanol” schedule to simulate a slow release of ethanol, and 18 mice received no injection. Death within 24 h of treatment qualified as lethality. One mouse died hours after a 4 mL/kg dose; however, the mouse was also under isoflurane anesthesia for a prolonged time (1 hour, 3%). Thus, considerations were made to limit isoflurane exposure after this adverse event. No other changes in systemic adverse events was noted for doses below 6 mL/kg. Mice were monitored for adverse events for a minimum of 14 days and up to 35 days (dictated by tumor growth) to observe any delayed effects after treatment. Greater than 200 breaths per minute qualified as respiratory distress. Limping or dragging of the tumor-bearing leg at any point indicated mobility impairment. Swelling and redness at any time after injection was noted as inflammation/edema. Bruising or bleeding of the tumor-bearing leg was noted as bleeding. Ulceration was defined as a scab on the skin over the tumor persisting for more than 3 days. A drop of 15% or greater of the baseline weight at any time after treatment was recorded as a severe loss in body weight. Because the tumors in this study were on the flank, mobility impairment, inflammation, edema, and ulceration of the tumor-bearing foot were considered localized adverse events. Lethality, respiratory distress, and loss in body weight were considered systemic adverse events.

### Quantification of Ethanol diffusion coefficient using Raman spectroscopy

The diffusion coefficient of ethanol through tumor tissue was measured using a Raman spectroscopy assay coupled with a deterministic mathematical model of ethanol diffusion. Tumor specimens (n=6) were cut longitudinally and trimmed to fit 12 mm Snapwell inserts containing porous polycarbonate membranes (Corning-Costar®, New York, NY). Each snap well was placed in a custom-built chamber filled with 600 µL of ethanol to enable ethanol diffusion through the membrane and upwards into the tumor (**Fig 2**). A custom-built confocal Raman spectroscope with a 785 nm excitation laser diode (LD785-SH300, Thorlabs Inc., Newton, NJ) was used to capture Raman spectra at the surface of the tumor (*30*). The least squares fit was performed in MATLAB (MathWorks, Natick, MA) to extract the relative contribution of ethanol and tumor tissue to the measured Raman spectra at the tumor surface, using a previously validated method (*30-32*).

To compute the diffusion coefficient of ethanol, a deterministic two-compartment transport model was used that recapitulates the Raman assay, using Fick’s law of diffusion to characterize ethanol diffusion from an ethanol compartment at the bottom, to the tumor at the top (**Supplemental Fig 1**). An analogous model was outlined previously in (*33*) and was applied to estimate the diffusion coefficients of drugs in biological tissues (*32*). Here, we assumed that the partition coefficient between the ethanol and tumor compartments (ΦET) was 1 due to the high porosity of the polycarbonate membrane, which is fully permeable to small molecules like ethanol. The diffusion coefficient of pure ethanol is 10^−5^ cm^2^/s (*34*), and the height of the ethanol compartment, hE, was 1.5 cm. The height of each tumor sample, hT, was measured using a micrometer. The two unknowns, the diffusion coefficient of ethanol in the tumor (DT) and the first-order loss term that accounts for ethanol evaporation from the tumor (KEV) were computed and optimized by fitting the predicted concentration of ethanol at the surface of the tumor to the measurements obtained using Raman spectroscopy.

**Figure 1:**
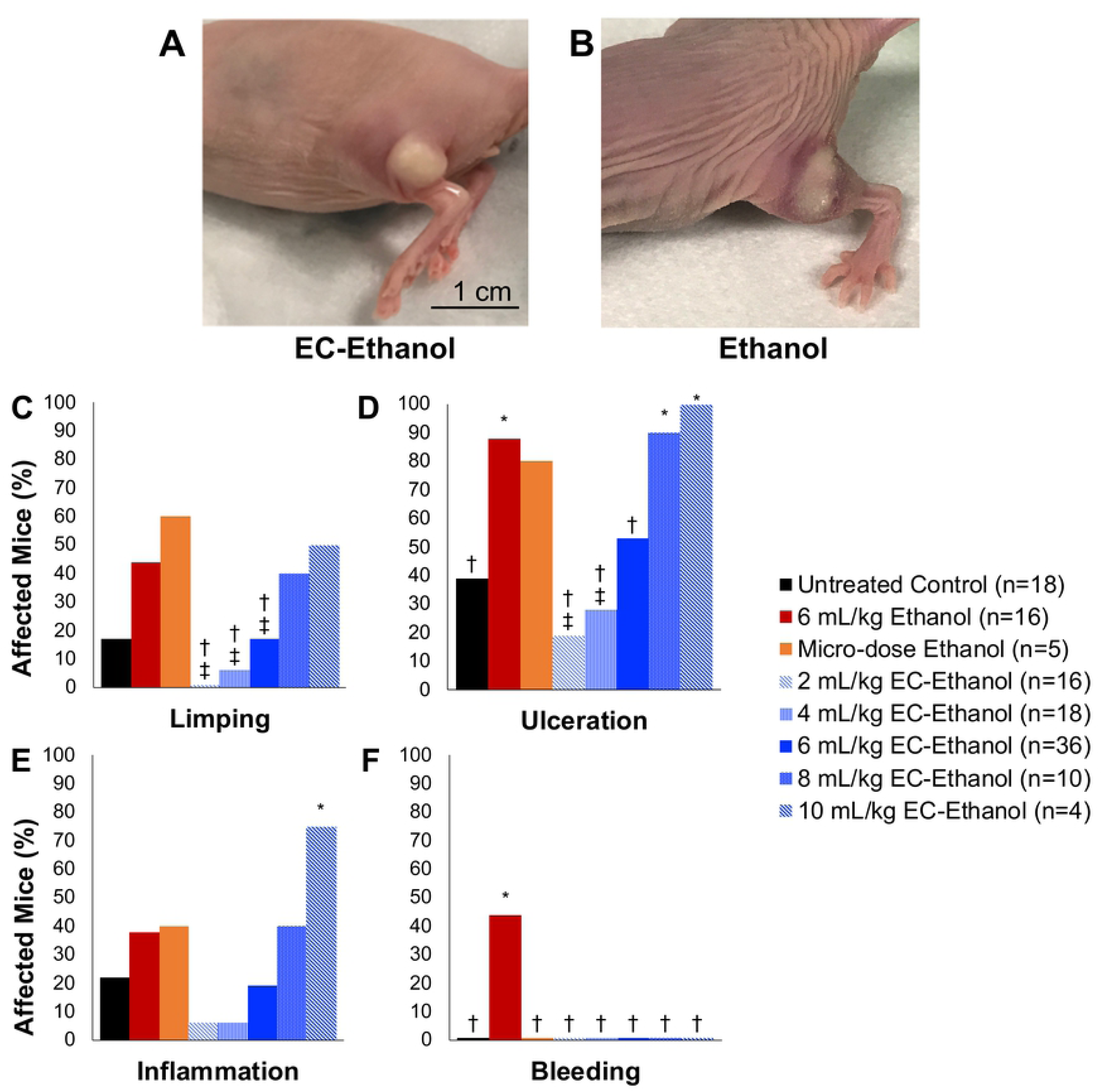
Localized Adverse Events after EC-ethanol Compared to Pure Ethanol. Mouse flank tumor after injection of (**A**) 6% EC-ethanol and (**B**) ethanol. Rates of (**C**) limping, (**D**) tumor ulceration, (**E**) inflammation, and (**F**) subdermal bleeding after each treatment protocol. Athymic nude mice with 67NR tumors grown on their flanks were injected at a rate of 1 mL/hr when tumors were approximately 50 mm^3^ in volume. Chi squared test significance of *P*<0.05 is indicated with (*) asterisk compared to untreated group, (†) dagger compared to 6 mL/kg ethanol group, (‡) double dagger compared to repeated 6 x 2 mL/kg/day ethanol group.

**Figure 2:**
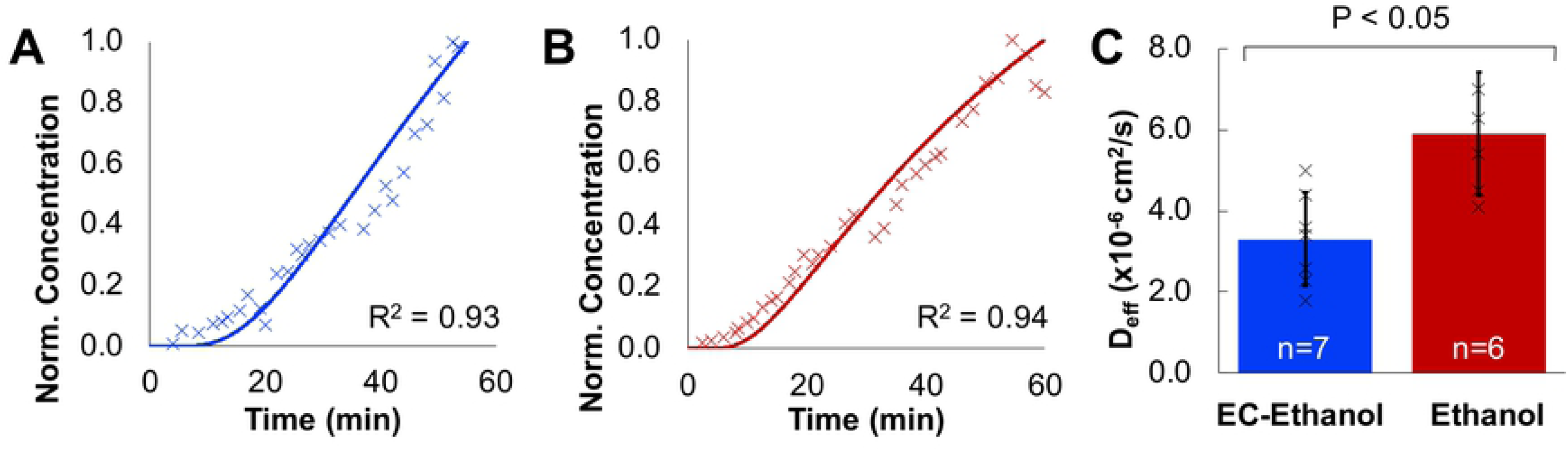
Quantification of Diffusion Coefficient of EC-Ethanol and Pure Ethanol with Raman Spectroscopy. (**A**) Representative ethanol concentration measurements for 6% EC-Ethanol measured with Raman spectroscopy and fit to a diffusion curve for D_eff_ of 1.9×10^−6^ cm^2^/s. (**B**) Representative ethanol concentration for pure ethanol measured with Raman spectroscopy and fit to a diffusion curve for D_eff_ of 5.4×10^−6^ cm^2^/s. (**C**) The effective diffusion coefficient was slower when using EC-ethanol (n=7) compared to pure ethanol (n=6). Bars indicate SD. \**P*<0.05 using Tukey’s HSD test.

### Whole-body Fluorescent Imaging

Mice were imaged using a whole-body *in vivo* IVIS Lumina imaging system (Perkin Elmer) in the prone position, 30 minutes after injection with 6% EC-ethanol or pure ethanol (n=5) to allow for injectate diffusion. Images were acquired with excitation at 430 nm, emission at 520 nm, and an exposure time of 1 second. Mice with 5 mm 67NR flank tumors (n=5) were given an injection of 6 mL/kg of either 6% EC-ethanol or pure ethanol. Pixels with a radiant efficiency of 0.5×10^10^ or higher we selected to represent the injection area. Area calculations and compactness calculations were performed using standard image processing tools in MATLAB (MathWorks).

### Imaging with a hand-held fluorescence Microscope

For *ex vivo* imaging, 6% EC-ethanol was injected into ∼1000 mm^3^ tumors. Tumors were injected with 150 μL of either pure ethanol (n=10), 3% EC-ethanol (n=10), or 6% EC-ethanol (n=20) at 1 mL/h. A control image of a tumor without any injection was acquired to account for tumor autofluorescence. Tumors were flash frozen and cross-sectioned centrally for imaging at a 10 mm working distance. Tumors were imaged using a fluorescence microscope with excitation at 480 nm and an emission bandpass filter at 520±15 nm. Pixels with >80% saturation in the green channel were counted as the area of fluorescence. Pixel dimensions were converted to an area using a digital ruler to scale each set of photos, and the radius r_1_ was computed. The fluorescent volume V_1_ (calculated using r_1_) was calculated assuming the fluorescent image was taken at the center of a spherically distributed injection.

### Histology and Immunohistochemistry

For viability staining, a 5 µm section was taken serially every 2 mm throughout the tumor and stained with NADH-diaphorase to quantify necrosis. NADH-diaphorase staining was performed using Nitrotetrazolium Blue Chloride (Sigma-Aldrich) and B-Nicotinamide Adenine Dinucleotide (Sigma-Aldrich). The region of interest (ROI) was selected manually from a digital scan at 5x magnification to include only non-viable cellular regions. Adjacent H&E slides were used to confirm tumor regions. Necrotic volume was calculated by the Reimann Sum: 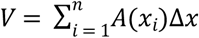, where volume (V) is equal to the summation of area (A) from 1 to n times the distance between each measurement (x). In this case, x = 2 mm.

### Tumor Growth and Survival

For tumor growth and survival studies, mice received 6% EC-ethanol injection four days after cell inoculation, when tumors were palpable and ∼50 µL in volume. Mice were assigned to receive either no treatment (n=8), a single injection of pure ethanol at 6 mL/kg (n=6), repeated injections of pure ethanol at 50 µL/day for 6 days (n=5), or a single injection of 6% EC-ethanol at 6 mL/kg (n=10). Mice were euthanized when the tumor reached 2000 mm^3^ (2 cm^3^). Time to a tumor volume of 1500 mm^3^ was used as a surrogate for survival time. If a humane endpoint was met before the maximum tumor burden, the humane endpoint was recorded as an adverse event. Tumor length (L) and width (W) was measured using calipers, and volume (V) was calculated using V = (W^2^ × L)/2. Average tumor volume did not include deceased mice.

### Statistical Analysis

Rates of adverse events were assessed with a Chi Squared test with a confidence level of 95% with a degree of freedom of 1. Two-tailed ANOVA testing with unequal variance was performed on all groups against each other and controls with a confidence level of 95%. Post-hoc multiple comparisons were performed using Tukey’s HSD test. Survival curves were quantified using Kaplan-Meier analysis, and a log-rank test was performed to determine the significance of a *P*-value less than 0.05 with a confidence level of 95%.

## Results

### EC-Ethanol Significantly Reduces Adverse Events compared to Pure Ethanol

We investigated whether using EC-ethanol would impact the rates of adverse effects compared to pure ethanol injections (recorded in **Supplemental Table 1)**. Mice with 67NR flank tumors received a single intra-tumoral injection of different volumes of 6% EC-ethanol (2-10 mL/kg), 4 days after tumor inoculation when tumors were approximately 50 μL in volume. A dose of 6 mL/kg of EC-ethanol was identified as the maximum tolerable dose (MTD) because it did not cause significantly more systemic adverse than untreated tumor bearing controls (**Supplementary Table 1**). These mice were compared to both mice that received a single intra-tumoral injection of 6 mL/kg (150 μL for average 25 g mouse) of ethanol or a repeat dose of 2 mL/kg (50 μL for average 25 g mouse) of ethanol over 6 days. All injections were delivered at a rate of 1 mL/hr. An untreated tumor bearing group was also included. A mouse was considered to have a localized adverse event if it displayed signs of mobility impairment (limping), subdermal bleeding, inflammation, or ulceration (scabbing) at any time during the two-week monitoring period (**Fig 1A-B**). Mobility impairment in the form of dragging the tumor-bearing leg or limping was most likely due to muscle or nerve damage in the leg from either ethanol or tumor invasion. EC-ethanol mixtures reduced limping, ulceration, and bleeding, but not inflammation, when compared to the same dose of pure ethanol (**Fig 1C-F**). A 6 mL/kg dose of EC-ethanol resulted in less local bleeding, inflammation, and limping than the same dose of pure ethanol, indicating that EC helps limit off target effects of intratumoral ethanol injections.

### EC-Ethanol Slows Ethanol Diffusion

In order to investigate the mechanism for improved safety of EC-ethanol over ethanol, the release kinetics of ethanol through tissue were investigated using Raman Spectroscopy. The tissue-depot interface was modeled as shown in **Supplemental Fig 1**. EC was hypothesized to reduce off-target leakage by slowing the release of ethanol from the injected depot site. The relative concentration of ethanol over time a sample was measured with Confocal Raman Spectroscopy (*30-32*) as ethanol diffused through thick (5 - 10 mm) sections of 67NR tumors. The change in ethanol concentration over time was used to fit a model of molecule transport in a two-compartment model. The effective diffusion coefficient (D_eff_) of a molecule represents the effective transport rate at which molecules move from one compartment to the other. D_eff_ can be approximated from the concentration over time recordings (**Fig 2A-B**) as in previously validated algorithms (*30-32*). The D_eff_ of ethanol of 6%EC-ethanol was approximately half that of pure ethanol (3.3 ± 1.5 × 10^−6^ cm^2^/s vs. 5.9 ± 1.2 × 10^−6^ cm^2^/s; *P*<0.05; n=7,6) (**Fig 2C**). Taken together, these results demonstrate that addition of EC to ethanol significantly slows the release of ethanol through tumor tissue.

### EC-Ethanol Delivery is More Uniformly Distributed Compared to Pure Ethanol

To demonstrate that the slower diffusion of EC-ethanol influences the local distribution of ethanol, the localization of EC-ethanol and ethanol-only injections were quantified. Both EC-ethanol and pure ethanol were mixed with the fluorescent dye, fluorescein (2.5% w/w) to enable *in vivo* tracking of the injectate. Small, 50 mm^3^ 67NR tumors were injected with 150 µL at 1 mL/hr to force overflow from the tumor and monitor the event of tumor leakage. Nude mice were used to limit autofluorescence from fur. Whole-body imaging was performed 30 min after injection (**Fig 3A, B)** using an *in vivo* imaging system (IVIS Lumina, Perkin Elmer. Compactness— defined as 4πA/P^2^, where A and P are the area and perimeter of the fluorescent region, respectively—was calculated using MATLAB. A perfect circle has a compactness of one, and any deviations from a circle will have a compactness that deviates from one. The compactness of the EC-ethanol injectate was significantly greater than that of pure ethanol (0.96 ± 0.01 vs. 0.80 ± 0.13; *P*<0.05; n=8,9) (**Fig 3C**), indicating reduced injectate leakage when using the EC-ethanol mixture over pure ethanol.

**Figure 3.**
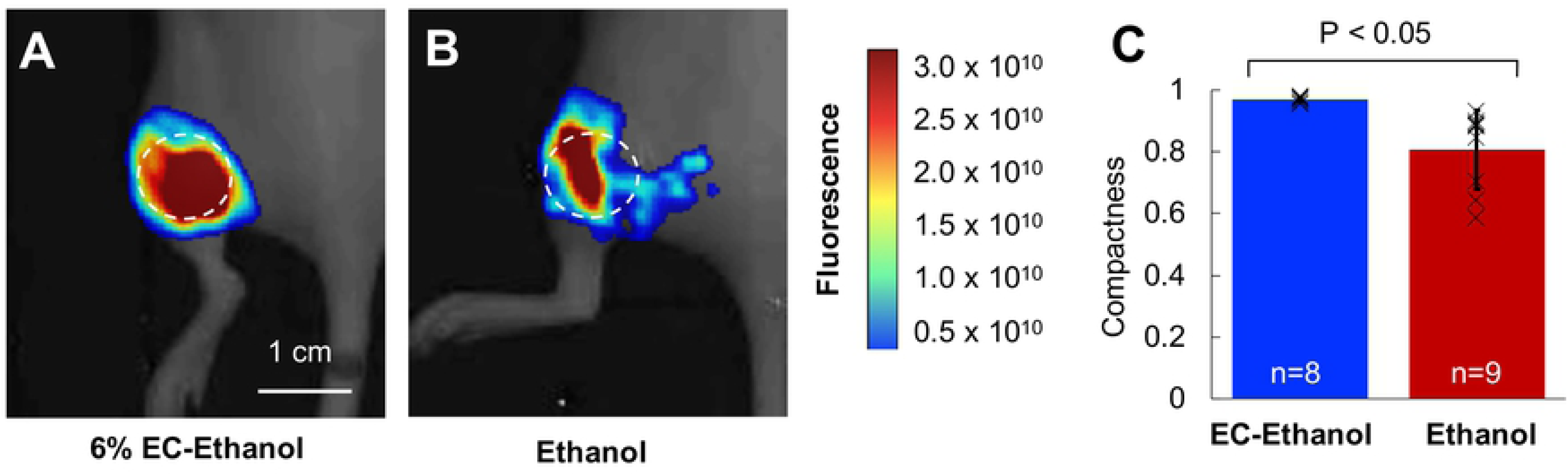
Injection of EC-ethanol Yields a More Compact Injection Zone than Pure Ethanol. (**A, B**) *In vivo* images showing the fluorescent volume following a single injection of 150 µL of 6%EC-ethanol or pure ethanol. Images were acquired 30 min after injection. The dashed circle indicates the approximate tumor location. **(C**) The compactness of the injectate was greater for EC-ethanol (n=8) than for pure ethanol (n=9).

### EC-ethanol Increases Intra-tumoral Ethanol Retention

The effect of relative concentration of EC and injection rate on injectate retention in the tumor were evaluated. A large, 1 cm^3^ tumor volume was selected to replicate clinically relevant tumor sizes while remaining under the ethical tumor burden limit for mice of 2 cm^3^. Nude mice with 1 cm^3^ 67NR flank tumors received a single 150 μL intra-tumoral injection of either pure ethanol, 3% EC-ethanol, or 6% EC-ethanol. Injections of pure ethanol (n=10), 3% EC-ethanol (n=10), and 6% EC-ethanol (n=20) were performed at 1 mL/hr Injections and also at 5 mL/hr (n=5) and 10 mL/hr (n=5) for 6% EC-ethanol injections. The contrast agent fluorescein (2.5% w/w) was added to image intra-tumoral injectate retention using fluorescence microscopy. The area of fluorescein was quantified in frozen tumor cross sections immediately after injections as outlined in the methods section. The intra-tumoral fluorescein retention was greater following injection of 6% EC-ethanol than following injection of 3% EC-ethanol or pure ethanol (*P*<0.01; all others n.s.) (**Fig 4A,B**). The intra-tumoral fluorescein retention was also greater when using slower injection flow rates: injection of 6% EC-ethanol at 1 mL/hr yielded a greater fluorescent volume than at 5 or 10 mL/hr (*P*<0.05) (**Fig 4A, B**). The ratio of fluorescent volume to injected volume was 1.85 ± 1.00 for 6% EC-ethanol, which was significantly greater than that for 3% EC-ethanol (1.02 ± 0.75, P<0.05) or pure ethanol (0.74 ± 0.58. P<0.05).

**Figure 4.**
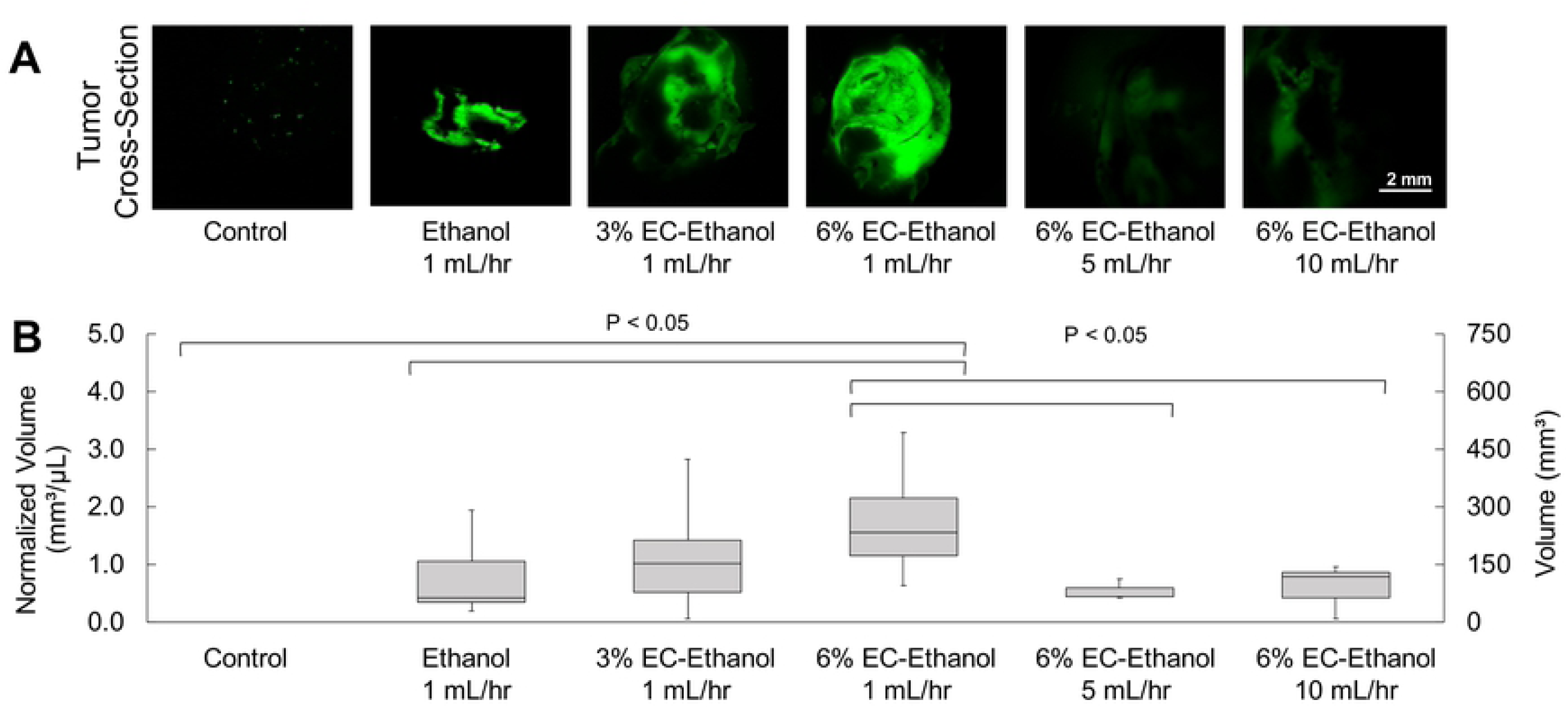
Relationship between EC concentration, flow rate and intra-tumoral ethanol retention. (**A**) Widefield fluorescence microscopy of large (1 cm^3^) 67NR flank tumors cross-sections after 150 μL injections of pure ethanol, 3% EC-ethanol, or 6% EC-ethanol at flow rate of 1 mL/hr and for 6% EC-ethanol at additional flow rates of 5 mL/hr and 10 mL/hr. (**B**) Intra-tumoral retention volume normalized by injection volume of 150 µL.

### 6% EC-ethanol Results in Larger Tumor Necrotic Volumes Compared to 3% EC-ethanol or Pure Ethanol

Next, viability staining was performed to confirm that fluorescein retention corresponded to ethanol exposure, and thus necrosis. 67NR tumors were selected because they experience minimal natural tumor necrosis at large volumes, as seen in the untreated mice here. The necrotic volume following EC-ethanol injections was assessed with NADH-diaphorase viability staining. A large (1 cm^3^) tumor volume was selected to replicate clinically relevant tumor sizes while remaining under the ethical tumor burden limit. After grown to approximately 1 cm^3^, 67NR tumors were injected with 150 μL of pure ethanol, 3% EC-ethanol, or 6% EC-ethanol at 1 mL/hr. At 24 hours post treatment, tumors were flash frozen, serially sectioned every 2 mm, and stained with Hematoxylin and Eosin (H&E) to confirm tumor presence (**Fig 5A**) and NADH-diaphorase to distinguish viable cells (blue) from necrotic cells (white) (**Fig 5B)**. Untreated tumors were included as a control. The necrotic volume following injection of 6% EC-ethanol was 5-fold greater than that of pure ethanol. The average ratio of necrotic volume to injected volume for 6% EC-ethanol was 2.00 ± 0.42, which was significantly greater than 3% EC-ethanol (0.80 ± 0.2) or pure ethanol (0.40 ± 0.09) (**Fig 5C**). As inducing necrosis is the primary objective of any ablation technique, 6% EC-ethanol is more effective at inducing target tissue necrosis compared to 3% EC-ethanol or pure ethanol.

**Figure 5:**
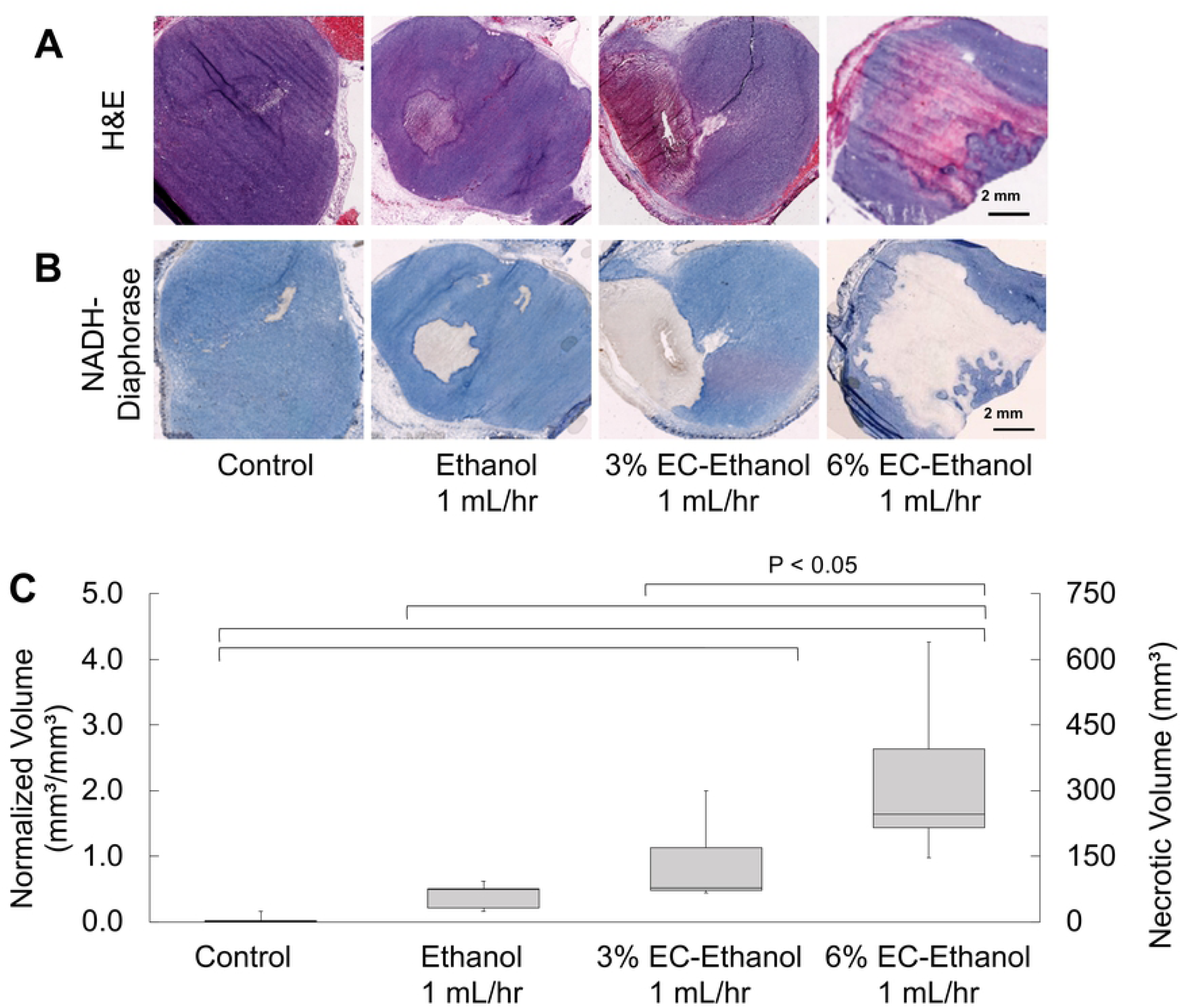
Relationship between EC-ethanol Concentration and Necrosis. Representative images of (**A**) H&E stained sections and (**B**) adjacent NADH-diaphorase stained sections from frozen blocks. (**C**) Ratio of necrotic volume to injected volume (normalized volume) after injection of 150 μL of pure ethanol at 1 mL/hr (n=5), 3% EC-ethanol at 1 mL/hr (n=5), or 6% EC-ethanol at 1 mL/hr (n=5). * *P*<0.05 using Tukey’s HSD test.

### EC-ethanol Reduces the Rate of Tumor Growth and Increases Survival Rates compared to Pure Ethanol, Obviating the Need for Repeat Injections

To quantify long-term therapeutic efficacy, 67NR tumors in BALB/c mice grown to approximately ∼50 mm^3^ in volume were randomly assigned to receive either no treatment (n=8), 150 µL of ethanol (n=6) or 150 µL of 6% EC-ethanol (n=6) at a rate of 1 mL/hr. Additionally, a repeated schedule of 50 µL was used for daily injections of ethanol for 6 days, to test the hypothesis that EC-ethanol is functionally similar to a slow release of repeated ethanol injections (n=6). Tumor size was measured with calipers three times per week. Time to a tumor volume of 1500 mm^3^ was used as a proxy for survival time. Average tumor growth rates are reported in **Figure 6A**. Survival probability was quantified with Kaplan-Meier curves in **Figure 6B**. Mice treated with a single ethanol injection did not experience a growth delay compared to untreated controls. Only mice treated with EC-ethanol experienced significantly increased survival rates compared to untreated controls and the same dose of ethanol (**Fig 6B**). Repeated ethanol injections did not produce significantly greater survival times compared to any group (**Fig 6B**).

**Figure 6.**
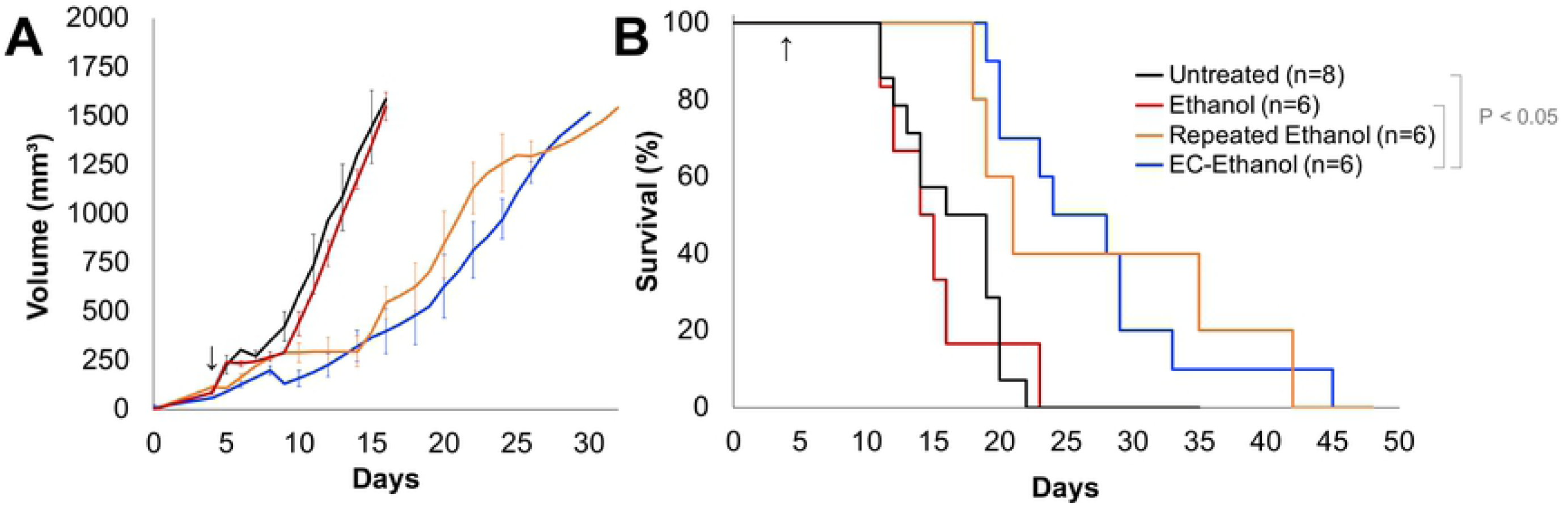
Increased Survival following 6% EC-ethanol Injections Compared to the Same Dose of Pure Ethanol: (**E**) Average tumor growth for all groups. (**F**) Kaplan-Meier survival probability. Arrow indicate first day of treatment. Log-rank test on Kaplan-Meier survival curves, pair-wise *P*<0.05, all other pairs n.s.

## Discussion

In this study it was demonstrated that 6% EC-ethanol delivered at a rate of 1 mL/hr is safer and more effective in treating 67NR tumors as compared to the same dose and rate of delivery of pure ethanol. Doses below 150 µL were found to be safe when injected in mice flank tumors. 6% EC slowed the effective diffusion of ethanol through tumor tissue. Intratumoral injections of 6% EC-ethanol produced a necrotic volume that is 5 times larger than the same dose of pure ethanol. Finally, it was demonstrated that a single dose of 6% EC-ethanol increased average survival times over the same dose of pure ethanol and untreated controls. Notably, survival rates of mice treated with 150 µL 6% EC-ethanol were not significantly different from six 50 µL repeated pure ethanol injections. We have shown here that a single EC-ethanol can limit tumor growth for aggressive murine 67NR tumors, where ethanol ablation was previously ineffective, requiring multiple injections to slow tumor growth.

While EC-ethanol increased tumor necrosis and survival, no mice lived longer than 45 days after tumor inoculation. There are several potential ways to improve the efficacy of EC-ethanol ablations to increase the number of mice that achieve complete cure. One straightforward approach is to inject in multiple regions throughout the tumor. In this study the needle was placed in the center of the tumor for the duration of the injection, however multiple spatially separated injections may further improve efficacy by providing complete tumor coverage which could be confirmed with ultrasound in larger tumor models. Because the maximum tolerable dose (MTD) given in this study was limited by the body weight of the mouse, larger dose-to-tumor volume ratios could be tested in larger animal models. In humans, larger injection volumes could be used, and the injection rate may need to be faster than 1 mL/hr to deliver large enough doses in a reasonable amount of time. Notably, a 150 µL injection of 6% EC-ethanol resulted in 300 mm^3^ of necrosis which was 5-fold greater than that of pure ethanol alone. Thus, 6% EC-ethanol may enable the ablation of larger lesions than what was previously possible with ethanol alone (∼ 2 cm) (*19*). While future studies are needed to compare EC-ethanol to standard-of-care treatments, EC-ethanol ablation may enable safe use of ethanol ablation for more indications than previously imagined. Further studies are needed to confirm the clinical feasibility of EC-ethanol for tumors in other locations.

Additionally, ablation with EC-ethanol injections could double as a delivery carrier for additional therapeutics as demonstrated here with fluorescein. More fluorescein was retained in the tumor when EC was included with ethanol injections, thus hydrophobic molecules with similar size and polarity to fluorescein such as paclitaxel, docetaxel, and tamoxifen may behave similarly in solution with EC-ethanol. Combining ablation with chemotherapy could potentially create a more robust response compared to either one alone (*35*). Additionally, immunomodulatory agents may enhance EC-ethanol therapy. As EC-ethanol creates a large necrotic region like other ablation modalities, there is a potential for adaptive immune system activation through the release of tumor antigens as seen with cryotherapy and thermal ablation (*36*). Moreover, the response to immunotherapies is known to be bolstered with local tumor ablation (*36*). Thus, future studies should investigate the effect that EC-ethanol ablation has on drug delivery and antitumor immunity.

In conclusion, it was demonstrated that augmenting percutaneous ethanol injections with the EC-ethanol treatment improves therapeutic efficacy and safety compared to injection of pure ethanol. The practical advantage of EC-ethanol ablation is in the reduction of number of treatment sessions and the reduced dose of ethanol needed to achieve the same efficacy. Further, EC-ethanol ablations are ultra-low-cost, allowing their use in low- and high-resource settings alike in the absence of bulky, expensive machinery. Thus, EC-ethanol may provide an additional line of treatment enabling more patients to access appropriate treatments.

## Acknowledgments

We thank M. Madonna, and T. Vincent for animal handling and tumor measurements. We acknowledge the Duke rodent breeding core for providing the nude mice; the Duke Substrate Core Services for pathology processing and NADH-diaphorase staining; the Duke Light Microscope Core Facility for microscopy imaging services; the Optical Molecular Imaging and Analysis for IVIS imaging services.

## Author Contributions

C.A.N. designed experiments and wrote the manuscript. C.A.N, R.M., E.C., D.A.A., I.B., B.T.C, collected data, analyzed data, prepared figures, and edited the manuscript. R.M. developed the ethanol-ethyl cellulose concept. C.T.L. developed the hand-held fluorescent microscope used in this study. Loren Baugh edited the manuscript text. A.A.S. assisted with interpretation of data in a clinical context, revising the manuscript for important intellectual content. B.T.C., J.L.M., D.K., M.W.D., J.I.E., N.R. provided study design guidance, analysis guidance, and reviewed the manuscript. J.I.E. provided pathology and animal model expertise. N.R. conceived the idea of fluorescent chemical ablation and guided the experimental design.

## Supporting Information Captions

**Supplemental Table 1. Counts of systemic and localized adverse events after ethanol injections**. Number of mice affected by local adverse events following a single intratumoral injection of EC-ethanol or pure ethanol.

**Supplementary Figure 1:**
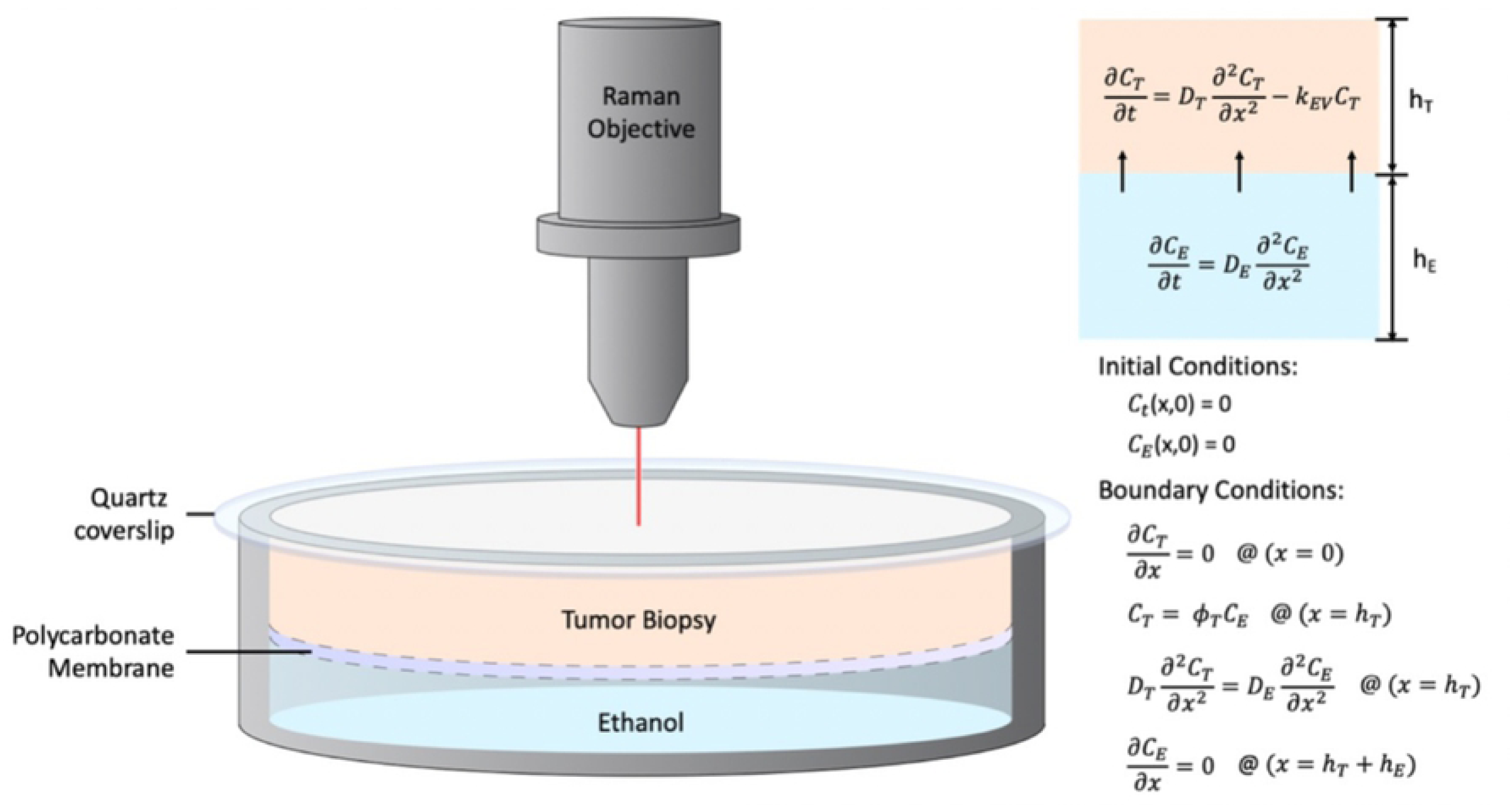
Raman Spectroscopy Diffusion Quantification. The methodology used to quantify the diffusion coefficient of ethanol in the tumor (DT). The experimental setup used for the Raman spectroscopy assay (left), consists of a tumor laying on a fully permeable membrane that allows seamless diffusion of ethanol upwards into the tumor. The CRS instrument captures the concentration of ethanol at the surface of the tumor over time. A deterministic transport model (right) characterizes ethanol transport and enables optimization of DT via fitting between the Raman measurements and the theoretical predictions.

## Notes

Funding: This study was supported by grants from the North Carolina Biotechnology Center Biotechnology Innovation Grant, the National Institutes of Health R21 (1R21CA241205-01), and the National Institutes of Health R01 (1R01CA239268-01).

The authors declare no potential conflicts of interest.

## References

1. A. N. Kurup, M. R. Callstrom, Ablation of skeletal metastases: current status. J Vasc Interv Radiol 21, S242–250 (2010).

2. A. N. Kurup, M. R. Callstrom, Image-guided percutaneous ablation of bone and soft tissue tumors. Semin Intervent Radiol 27, 276–284 (2010).

3. T. W. I. Clark, M. C. Soulen, Chemical Ablation of Hepatocellular Carcinoma. Journal of Vascular and Interventional Radiology 13, S245–S252 (2002).

4. M. D. Tito Livraghi et al., Long Term Results of Single Session Percutaneous Ethanol Injection in Patients with Large Hepatocellular Carcinoma. Cancer 83, 49–57 (1996).

5. A. D. S. A. Giorgio, G. De Stefano, U. Scognamiglio, N. Farella, A. Mariniello, V. Esposito, C. Coppola and V. Giorgio, Percutaneous Radiofrequency Ablation of Hepatocellular Carcinoma Compared to Percutaneous Ethanol Injection in Treatment of Cirrhotic Patients: An Italian Randomized Controlled Trial. Anticancer Research 31, (2011).

6. Y. J. Kim et al., Cystic versus predominantly cystic thyroid nodules: efficacy of ethanol ablation and analysis of related factors. Eur Radiol 22, 1573–1578 (2012).

7. J. H. Lee et al., Radiofrequency ablation (RFA) of benign thyroid nodules in patients with incompletely resolved clinical problems after ethanol ablation (EA). World J Surg 34, 1488–1493 (2010).

8. N. M. Lakkis et al., Echocardiography-Guided Ethanol Septal Reduction for Hypertrophic Obstructive Cardiomyopathy. Circulation 98, 1750–1755 (2001).

9. S. F. Nagueh et al., Comparison of ethanol septal reduction therapy with surgical myectomy for the treatment of hypertrophic obstructive cardiomyopathy. Journal of the American College of Cardiology 38, (2001).

10. M. Afshin Gangi et al., Interventional radiologic procedures with CT guidance in Cancer Pain Management. RadioGraphics 16, 1289–1304 (1996).

11. D. K. Filippiadis, S. Tutton, A. Mazioti, A. Kelekis, Percutaneous image-guided ablation of bone and soft tissue tumours: a review of available techniques and protective measures. Insights Imaging 5, 339–346 (2014).

12. L. M. Kenny, F. Orsi, A. Adam, Interventional radiology in breast cancer. Breast 35, 98–103 (2017).

13. F. D. K., S. Tutton, A. Kelekis, Percutaneous bone lesion ablation. La radiologia medica 119, 462–469 (2014).

14. M. Chin, C. L. Chen, K. Chang, J. Lee, J. Samarasena, Ethanol Ablation of a Peripheral Nerve Sheath Tumor Presenting as a Small Bowel Obstruction. ACG Case Rep J 3, 31–32 (2015).

15. D. H. Park et al., Endoscopic ultrasonography-guided ethanol ablation for small pancreatic neuroendocrine tumors: results of a pilot study. Clin Endosc 48, 158–164 (2015).

16. Y. N. Hasegawa S, Hiwaki T, Sako K, Komorizono Y, Baba Y, Imamura Y, Kubozono O, Yoshida A, Arima T, Factors that predict intrahepatic recurrence of hepatocellular carcinoma in 81 patients initially treated by percutaneous ethanol injection. Cancer: Interdisciplinary International Journal of the American Cancer Society. 86, 1682–1690. (1999).

17. M. Y. Koda M, Mitsuda A, Ohyama K, Horie Y, Suou T, Kawasaki H, Ikawa S, Predictive factors for intrahepatic recurrence after percutaneous ethanol injection therapy for small hepatocellular carcinoma.. Cancer 88, 529–537 (2000).

18. A. Facciorusso, G. Serviddio, N. Muscatiello, Local ablative treatments for hepatocellular carcinoma: An updated review. World J Gastrointest Pharmacol Ther 7, 477–489 (2016).

19. M. P. Giacomo Germani, Kurinchi Gurusamy, Tim Meyer, Graziella Isgrò, Andrew Kenneth Burroughs, Clinical outcomes of radiofrequency ablation, percutaneous alcohol and acetic acid injection for hepatocelullar carcinoma: A meta-analysis. Journal of Hepatology 52, 380–388 (2010).

20. E. H. Zubizarreta, E. Fidarova, B. Healy, E. Rosenblatt, Need for radiotherapy in low and middle income countries - the silent crisis continues. Clin Oncol (R Coll Radiol) 27, 107–114 (2015).

21. D. H. Robinson, Ethyl Cellulose-Solvent Phase Relationships Relevant to Coacervation Microencapsulation Processes. Drug Development and Industrial Pharmacy 15, 2597–2620 (1989).

22. R. Morhard et al., Development of enhanced ethanol ablation as an alternative to surgery in treatment of superficial solid tumors. Sci Rep 7, 8750 (2017).

23. R. Morhard et al., Understanding factors governing distribution volume of ethyl cellulose-ethanol to optimize ablative therapy in the liver. IEEE Trans Biomed Eng, (2019).

24. M. Schumacher et al., Treatment of venous malformations: first experience with a new sclerosing agent--a multicenter study. Eur J Radiol 80, e366–372 (2011).

25. A. Dompmartin et al., Radio-opaque ethylcellulose-ethanol is a safe and efficient sclerosing agent for venous malformations. Eur Radiol 21, 2647–2656 (2011).

26. M. Bellini et al., Percutaneous injection of radiopaque gelified ethanol for the treatment of lumbar and cervical intervertebral disk herniations: experience and clinical outcome in 80 patients. AJNR Am J Neuroradiol 36, 600–605 (2015).

27. C. Sanford, B. Mantooth, B. Jones, Determination of ethanol in alcohol samples using a modular Raman spectrometer.. Journal of Chemical Education. 78, (2001).

28. F. Li et al., Study of hydrogen bonding in ethanol-water binary solutions by Raman spectroscopy. Spectrochimica Acta Part A: Molecular and Biomolecular Spectroscopy. 189, 621–624 (2018).

29. L. Sm, R. Sz, G. Av, P. Ev, A. Cv, Thermal degradation of biodegradable blends of polyethylene with cellulose and ethylcellulose.. Thermochimica acta. 521, 66–73 (2011).

30. J. R. Maher et al., Co-localized confocal Raman spectroscopy and optical coherence tomography (CRS-OCT) for depth-resolved analyte detection in tissue. Biomed Opt Express 6, 2022–2035 (2015).

31. O. Chuchuen et al., Label-free analysis of tenofovir delivery to vaginal tissue using co-registered confocal Raman spectroscopy and optical coherence tomography. PLoS One 12, e0185633 (2017).

32. O. Chuchuen et al., Label-Free Measurements of Tenofovir Diffusion Coefficients in a Microbicide Gel Using Raman Spectroscopy. J Pharm Sci 106, 639–644 (2017).

33. Y. Gao, D. F. Katz, Multicompartmental pharmacokinetic model of tenofovir delivery by a vaginal gel. PLoS One 8, e74404 (2013).

34. C. J. Zhang, X. N. Yang, Molecular dynamics simulation of ethanol/water mixtures for structure and diffusion properties. Fluid Phase Equilibr 231, 1–10 (2005).

35. H. Takaki et al., Thermal ablation and immunomodulation: From preclinical experiments to clinical trials. Diagn Interv Imaging 98, 651–659 (2017).

36. R. Slovak, J. M. Ludwig, S. N. Gettinger, R. S. Herbst, H. S. Kim, Immuno-thermal ablations - boosting the anticancer immune response. J Immunother Cancer 5, 78 (2017).

